# There and back again: metagenome-assembled genomes provide new insights into two thermal pools in Kamchatka, Russia

**DOI:** 10.1101/392308

**Authors:** Laetitia G. E. Wilkins, Cassandra L. Ettinger, Guillaume Jospin, Jonathan A. Eisen

**Affiliations:** Department of Environmental Sciences, Policy &Management, University of California, Berkeley, CA USA; Genome Center, University of California, Davis, CA USA; Department of Evolution and Ecology, University of California, Davis, CA USA; Department of Medical Microbiology and Immunology, University of California, Davis, CA USA

**Keywords:** Archaea, hydrothermal, Kamchatka, metagenomics, metagenome-assembled genomes, terrestrial hot springs, tree of life, Uzon Caldera

## Abstract

Culture-independent methods have contributed substantially to our understanding of global microbial diversity. Recently developed algorithms to construct whole genomes from environmental samples have further refined, corrected and revolutionized the tree of life. Here, we assembled draft metagenome-assembled genomes (MAGs) from environmental DNA extracted from two hot springs within an active volcanic ecosystem on the Kamchatka peninsula, Russia. This hydrothermal system has been intensively studied previously with regard to geochemistry, chemoautotrophy, microbial isolation, and microbial diversity. Using a shotgun metagenomics approach, we assembled population-level genomes of bacteria and archaea from two pools using DNA that had previously been characterized via 16S rRNA gene clone libraries. We recovered 36 MAGs, 29 of medium to high quality, and placed them in the context of the current microbial tree of life. We highlight MAGs representing previously underrepresented archaeal phyla (*Korarchaeota, Bathyarchaeota* and *Aciduliprofundum*) and one potentially new species within the bacterial genus *Sulfurihydrogenibium*. Putative functions in both pools were compared and are discussed in the context of their diverging geochemistry. This study can be considered complementary to foregoing studies in the same ecosystem as it adds more comprehensive information about phylogenetic diversity and functional potential within this highly selective habitat.

## Introduction

Terrestrial hydrothermal systems are of great interest to the general public and to scientists alike due to their unique and extreme conditions. Hot springs have been sought out by geochemists, astrobiologists and microbiologists around the globe who are interested in their chemical properties, which provide a strong selective pressure on local microorganisms. Drivers of microbial community composition in these springs include temperature, pH, *in-situ* chemistry, and biogeography ^1–3^. The heated water streams contain substantial concentrations of carbon dioxide, nitrogen, hydrogen, and hydrogen sulphide. Moreover, high temperature subterranean erosion processes can result in high levels of soluble metals and metalloids. Microbial communities have not only developed strategies to resist these limiting conditions but have also invented ways to thrive by converting hot spring chemicals into energy ^4^.

The Uzon Caldera is part of the Pacific Ring of Fire and is one of the largest active volcanic ecosystems in the world ^4^. The geochemical properties of this system have been studied in detail ^5–7^. The systematic study of Kamchatka thermophilic microbial communities was initiated by Georgy Zavarzin in the early 1980’s ^8^. Briefly, the Uzon Caldera was created by a volcanic eruption. It is characterized by high water temperatures (20 – 95° Celsius), a wide range of pH (3.1 – 9.8), and many small lakes that are filled with sediment and pumice, dacite extrusions, and peatbog deposits ^9,10^. Most of the hydrothermal springs lack dissolved oxygen, but harbour sulphides and rare trace elements (antimony, arsenic, boron, copper, lithium, and mercury ^11^). Many previously undiscovered bacteria have been isolated in this region using culture-dependent methods ^12–14^, and 16S rRNA gene amplicon sequencing has revealed a diverse collection of new lineages of archaea ^15,16^. Although microorganisms from Uzon 17 Caldera are well represented in culture collections^17^, this region has remained relatively under-sampled by culture-independent methods ^8,9,11,18–23^.

In this study, we focus on two hydrothermal pools, Arkashin Schurf and Zavarzin Spring in the Uzon Caldera that were previously characterized using 16S ribosomal RNA gene sequencing and geochemical analysis by Burgess *et al*. ^9^ (Fig. 1). Arkashin Schurf (ARK) is an artificial pool, approximately 1 m^2^. in size, in the central sector of the East Thermal Field (54°30’0” N, 160°0’20” E), which was dug during a prospecting expedition 10 to Uzon by Arkadiy Loginov ARK has been generally stable in size and shape since its creation ^24^. Flocs ranging in colour from pale yellow-orange to bright orange-red have been observed floating in ARK ^10^. This pool is characterized by high concentrations of arsenic and sulphur, which result from the oxidation and cooling of magmatic waters as they reach the surface of the caldera ^9^. Zavarzin Spring (ZAV) is a natural pool, approximately 10 m^2^. in size, in the Eastern Thermal Field (54°29’53” N, 160°0’52” E). Unlike ARK, the size and shape of ZAV is constantly in flux as vents collapse and emerge and as the amount of snowmelt changes ^24^. Green microbial mats have been observed around the edge of ZAV and thicker brown and green mats have been found within the pool itself ^10^.

**Figure 1.**
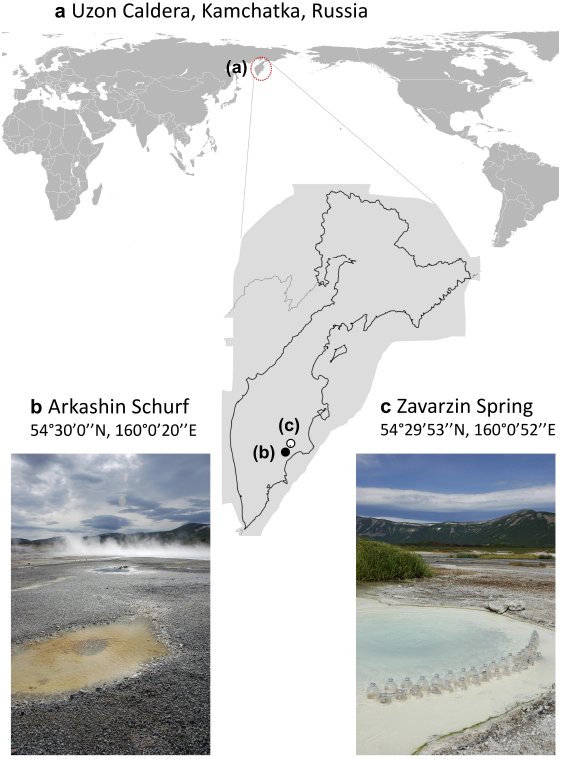
Sampling locations in the Uzon Caldera, Kamchatka Russia. DNA had been extracted in 2009 by Burgess *et al*. ^9^. from sediment samples of two active thermal pools Arkashin Schurf (b) and Zavarzin Spring (c). Photos were taken by Dr. Russell Neches during an expedition in 2012. Maps were plotted in Rv. 3.4.0 with the package ‘ggmap’ v. 2.6.1 ^97^

Burgess *et al*. ^9^ found that the two pools differed geochemically with ARK containing higher amounts of total arsenic, rubidium, calcium, and caesium; and ZAV containing higher amounts of total vanadium, manganese, copper, zinc, strontium, barium, iron and sulphur ^10^. Water temperatures in ARK ranged from 65°C near the vent to 32°C at the edge of the pool with temperatures as high as 99°C at 10 cm depth into the vent sediments. ZAV showed relatively lower temperatures between 26°C and 74°C at different locations of the pool.

Using the same DNA that had been used in the Burgess *et al*. study ^9^ as our starting material, we applied a metagenomic whole genome shotgun sequencing approach. We binned reads from the environment into individual population-specific genomes and then identified and annotated taxonomic and functional genes for the microorganisms in the two pools. In contrast to the 16S rRNA gene amplicon approach, metagenomic sequencing avoids taxonomic primer bias ^25^, provides more direct functional prediction information about the system ^26^, and ultimately can result in a more precise taxonomy through multi-gene and whole-genome phylogenetic approaches ^27^. However, at low sequencing depths, metagenomic sequencing and whole genome binning capture only the most abundant bacterial genes in the pools. Accordingly, we tested the following questions: (1) Can we recover metagenome-assembled genomes (MAGs) from the two pools? Are there any previously undiscovered or unobserved taxa that can be described using this approach? (2) How do any identified MAGs compare to Burgess *et al.’s* survey of the microbes in these pools? Do we find any archaea in the ARK pool from which Burgess *et al*. were unable to amplify any 16S rRNA gene sequences? (3) How do any MAGs found here fit into the current microbial tree of life? (4) Can we identify any differences in the functional genes or specific MAGs between the two pools that might be explained by their diverging geochemistry?

## Results

### Quality filtering and assembly

DNA libraries were prepared and then sequenced using Solexa3 84 bp paired-end sequencing. For ARK, 52,908,626 Solexa reads (4,444,324,584 bases) were processed while 77% of the reads were retained after adaptor removal and 76.32% passed trimming to Q10 (Supplementary Table S1). For ZAV, 58,334,696 Solexa reads were processed (4,900,114,464 bases) while 59.91% were retained and 59.05% passed the cut-offs. Reads for each sample were then assembled using SPAdes ^28^. The ARK assembly generated 103,026 contigs of sizes from 56 to 103,908 bp with an N50 of 3,059. The ZAV assembly generated 151,500 contigs of sizes from 56 to 791,131 base pairs with an N50 of 2637 (Supplementary Table S1). Sanger metagenomic reads were also generated from clone libraries for ARK and ZAV, but not used for metagenomic assemblies and binning.

### Metagenome-assembled genome quality and taxonomic identification

Using anvi’o ^29^, we assembled 36 draft MAGs, 20 from ZAV (three high-quality, 12 medium-quality and five low-quality) and 16 from ARK (seven high-quality, seven medium-quality and two low-quality; Table 1, Supplementary Table S2). These MAGs include only 11.7% of the nucleotides present in the ZAV assembly and 19.2% of the nucleotides in the ARK assembly. MAGs from ZAV and ARK were taxonomically inferred to be bacteria (n = 22; Fig. 2, Supplementary Fig. S1) and archaea (n = 14). MAGs from ARK were taxonomically assigned to a diverse group of 12 phyla (Table 2). A similar range in taxonomic diversity, 13 phyla, is seen in MAGs binned from ZAV (Table 3).

**Figure 2.**
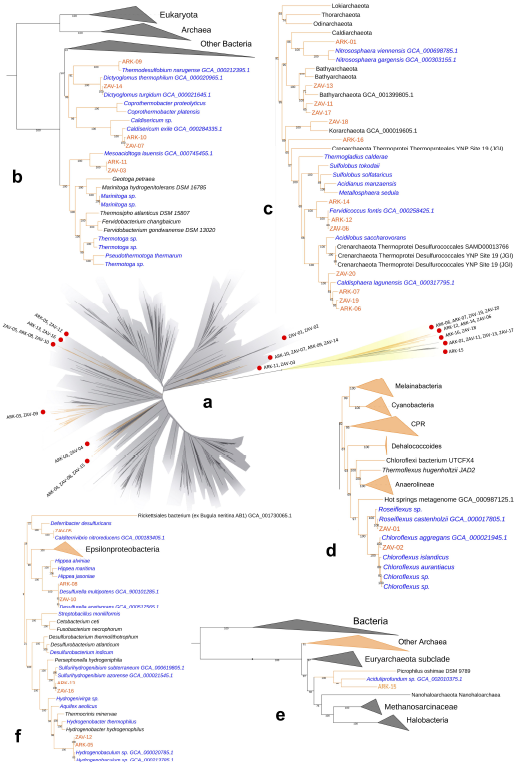
Placement of our MAGs into their phylogenetic context. Taxonomy of our MAGs (metagenome-assembled genomes) was refined by placing them into a phylogenetic tree using PhyloSift v. 1.0.1 with its updated markers database for the alignment and RAxML v. 8.2.10 on the CIPRES web server for the tree inference. This tree includes our 36 MAGs (red dots), all taxa previously identified by ^Burgess *et al*. (2012)^. with complete genomes available on NCBI (n = 148; ^80^), and 3,102 archaeal (yellow) and bacterial (grey) genomes previously used in Hug *et al*. (2016; ^77–79^). The complete tree in Newick format and its alignment of 37 concatenated marker genes can be found on Figshare ^73,96^. Branches with MAGs found in Arkashin Schurf (ARK) and Zavarzin Spring (ZAV) are enlarged (orange nodes). Blue: taxa from ^Burgess *et al*. (2012)^, black: taxa from Hug *et al*. (2016). GCA IDs from NCBI are shown for the closest neighbours of our MAGs. a) Current tree of life, reconstructed from Burgess *et al.;* b) Dictyoglomales, Thermoanaerobacteriales, Caldisericales, and Mesoaciditogales; c) Nitrosphaera, Bathyarchaeota, Korarchaeota, and Crenarchaeota; d) Chloroflexales; e) Euryarchaeota; and f) Deferribacteriales, Desulfobacteriales, and Aquificales. ARK-02, ARK-03, ARK-04, ZAV-04, ZAV-08, ZAV-09, and ZAV-15 can be found in Supplementary Figure S1.

**Table 1.**
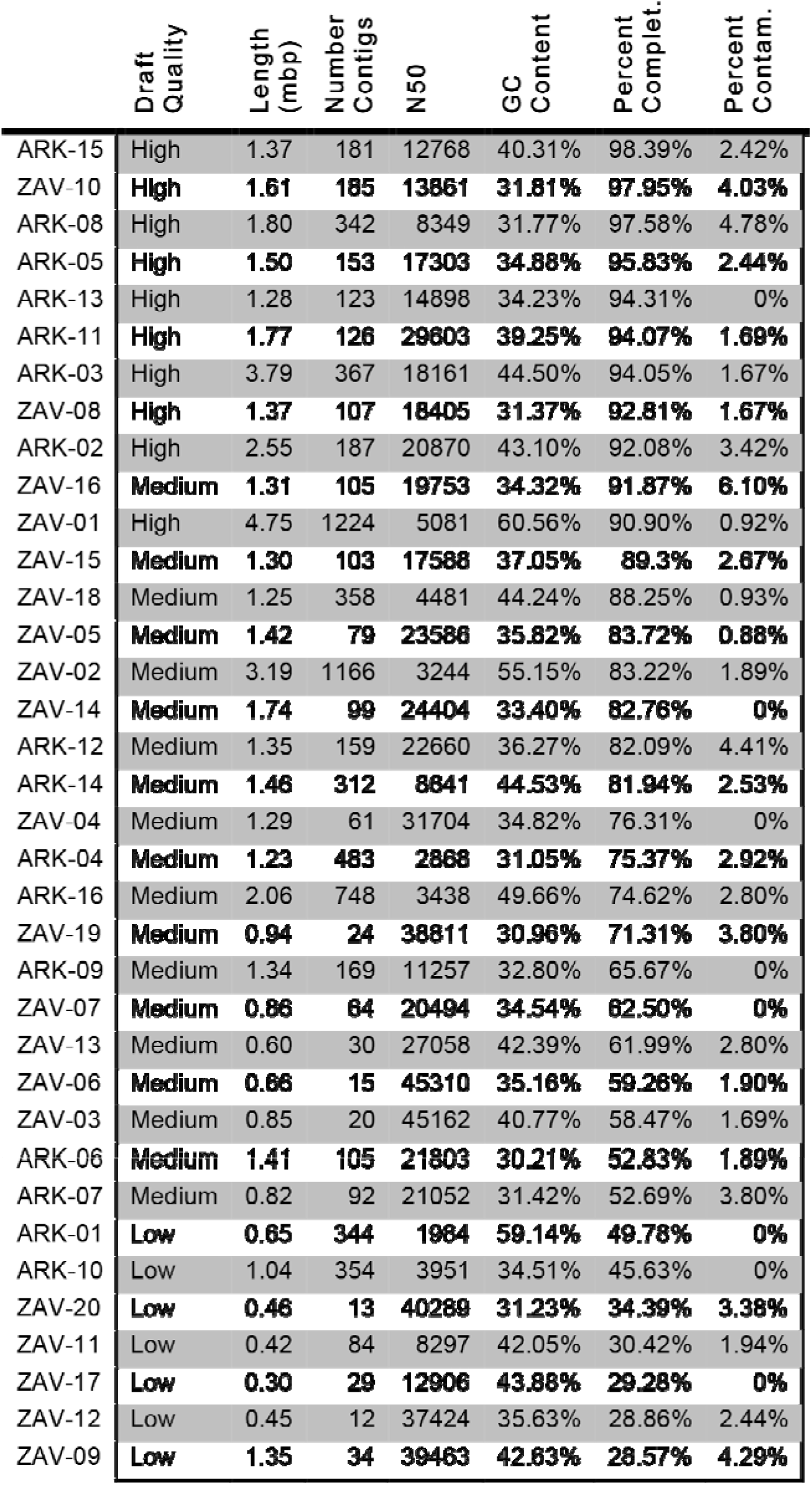
Genomic feature summary for metagenome-assembled genomes identified in Arkashin Schurf (ARK) and Zavarzin Spring (ZAV). Genomic features are summarized below for each metagenome-assembled genome (MAG) including length (mbp), number of contigs, N50, percent GC content, and completion and contamination estimates as generatedby CheckM. MAGs are sorted by percent completion and their draft-quality is indicated.

**Table 2.**
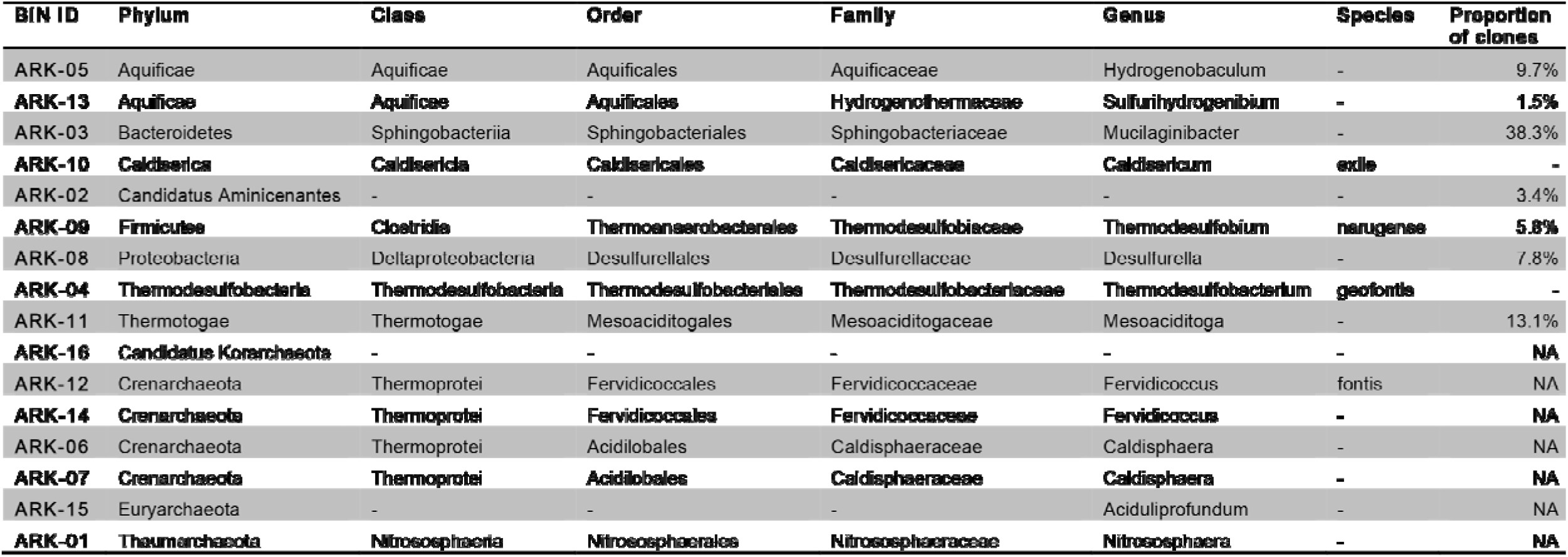
Taxonomic identification of MAGs in ARK. Here we report putative taxonomies for metagenome-assembledgenomes (MAGs) identified in Arkashin Schurf (ARK) and indicate their relative abundance in the reanalysed bacterial clone libraries constructed in Burgess *et al*. ^9^. They were unable to amplify archaeal sequences from ARK which is indicated in this table using ‘NA’.

**Table 3.**
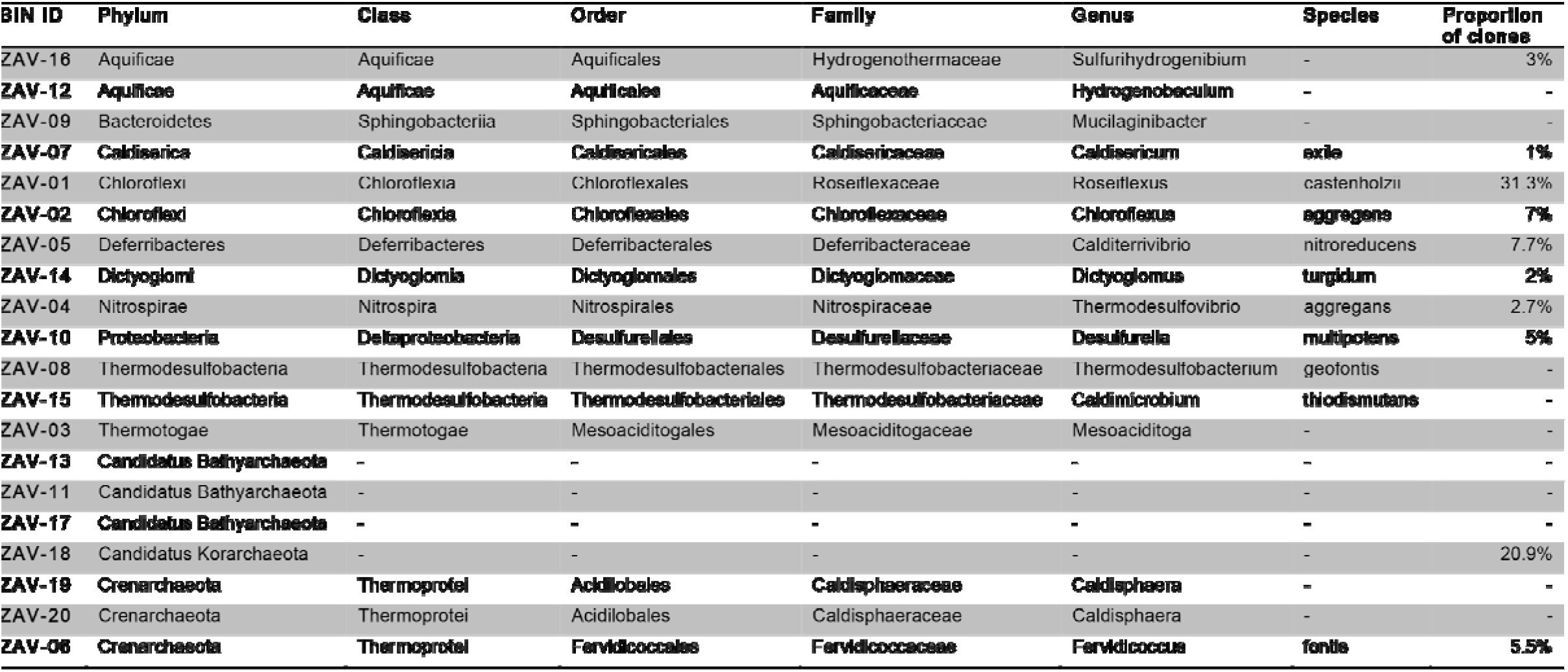
Taxonomic identification of MAGs in ZAV. Here we report the putative taxonomies for metagenome-assembled genomes (MAGs) identified in Zavarzin Spring (ZAV) and indicate their relative abundance in the reanalysed bacterial and archaeal clone libraries constructed in Burgess *et al*. ^9^.

### Phylogenetic placement of MAGs

Phylogenetic analysis placed MAGs into the following bacterial clades: two in the Chloroflexales (ZAV-01, ZAV-02; Fig. 2), one in Deferribacteriales (ZAV-05), two in Desulfobacteriales (ARK-08, ZAV-10), four in Aquificales (ARK-05, ARK-13, ZAV-12, ZAV-16), one in Dictyoglomales (ZAV-14), one in Thermoanaerobacteriales (ARK-09), two in Caldisericales (ARK-10, ZAV-07), two in Mesoaciditogales (ARK-11, ZAV-03), three in Thermodesulfobacteria (ARK-04, ZAV-08, ZAV-15; Supplementary Fig. S1), two in Sphingobacteriales (ARK-03, ZAV-09), one in Acidobacteriales (ARK-02), and one in Thermodesulfovibrio and sister to Dadabacteria (ZAV-04). Within the archaea, phylogenetic analysis placed MAGs into one of the following groups or positions: one MAG in Candidatus Nitrosphaera (ARK-01; Fig. 2), three in Bathyarchaeota (ZAV-11, ZAV-13, ZAV-17), one in Korarchaeota (ZAV-18), one sister to Crenarchaeota (ARK-16), and seven in Crenarchaeota (ARK-12, ARK-14, and ZAV-06 most closely related to *Fervidicoccus;* and ARK-06, ARK-07, ZAV-19, and ZAV-20 most closely related to *Caldisphaera)*. ARK-15 shared a common ancestor with an *Aciduliprofundum* species, nested within Thermoplasmatales and sister to Euryarchaeota. Compared to identification with CheckM there were two ambiguities: ARK-16 was assigned to Korarchaeota (Supplementary Table S3) vs. a sister to Crenarchaeota (Fig. 2); and ARK-02 was assigned to Candidatus Aminicenantes (Supplementary Table S3) *vs*. Acidobacteriales (Supplementary Fig. S1).

### Taxonomic inference of 16S rRNA gene sequences from Burgess et al.

Burgess *et al*. ^9^ previously generated 16S rRNA gene sequences from clone libraries to investigate the archaeal and bacterial diversity of these pools. By downloading the Burgess *et al*. ^9^ 16S rRNA gene sequence data and analysing it using an updated database, we were able to infer taxonomy for some sequences that were previously unclassified (Tables S4, S5, S6). We identified representatives of two new archaeal phyla (Aenigmarchaeota and Thaumarchaeota) in ZAV and saw a decrease in the proportion of unidentified archaeal sequences from the 13% reported by Burgess *et al*. to 7.7%. We also found no representatives of Euryarchaeota, which had previously been reported as 7% of the sequence library and suspect that these reads may have been reassigned to different taxonomic groups due to updates to the Ribosomal Database Project (RDP) database. Several previously unobserved phyla were identified as small proportions of the ZAV bacterial sequence library including Actinobacteria (0.3%), Atribacteria (4%), Elusimicrobia (1%), Ignavibacteriales (2.7%) and Microgenomates (0.3%). We saw only a moderate decrease in the unclassified bacteria from 24% to 17.3%. In comparison, we report a large decrease in the proportion of unclassified bacteria in the ARK sequence library from 19% to only 2.9%. This can be attributed to the identification of two previously unobserved phyla in the ARK bacterial library, Candidatus Aminicenantes (3.4%) and Thermotogae (13.1%).

### Taxonomic comparison to 16S rRNA gene sequence libraries

Taxonomic assignments for seven of the nine bacterial MAGs found in ARK placed them in the same genera identified from the clone libraries prepared by Burgess *et al*. ^9^ (Table 2). Since Burgess *et al*. were unable to amplify archaeal sequences from ARK, there were no 16S rRNA gene results on archaea to which to compare. Here, we were able to identify archaea from four different archaeal phyla including representatives of novel lineages. Eight of the thirteen bacterial and two of the seven archaeal MAGs found in ZAV match genera from the libraries constructed in Burgess *et al*. ^9^ (Table 3).

Further comparing the 16S rRNA gene sequence libraries from ARK and ZAV to the inferred phyla present in the Sanger metagenomes prepared by TIGR and the quality-filtered Solexa reads, we found that the latter were able to detect additional phyla present in these hydrothermal systems (Tables S7, S8). All of the assembled MAGs matched phyla observed using either all three methods (16S rRNA gene sequence libraries, Sanger metagenomes, Solexa reads) or using both the Sanger metagenomes and Solexa reads, but not the 16S rRNA gene sequence libraries (Fig. 3).

**Figure 3.**
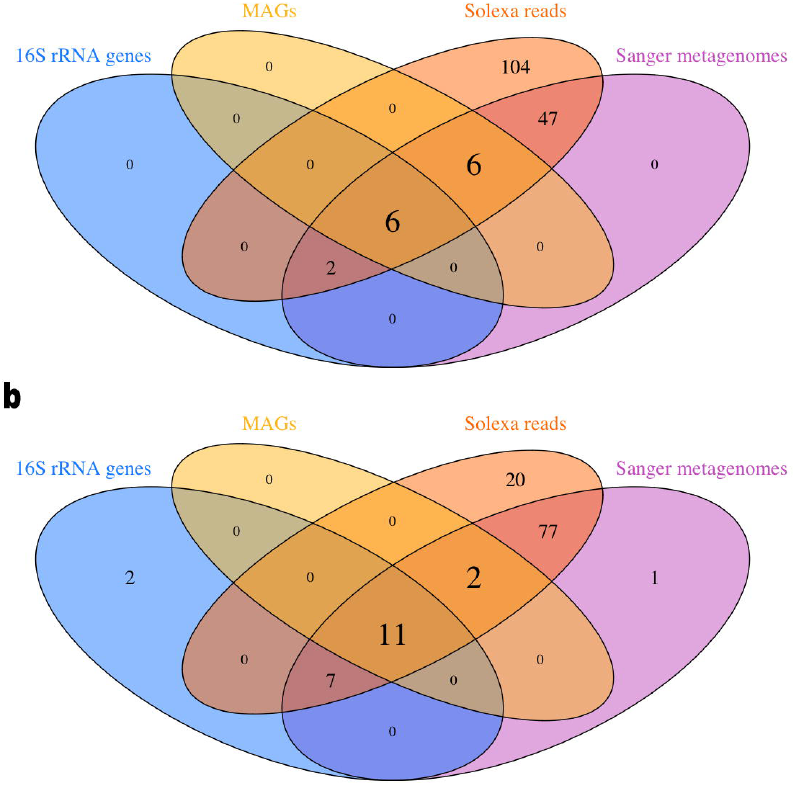
Shared phyla between MAGs and different sequencing methods. Venn diagrams depict the number of shared phyla observed between metagenome assembled genomes (MAGs) and different methods of sequencing and taxonomic assignment for (a) Arkashin Schurf (ARK) and (b) Zavarzin Spring (ZAV). The different methods include the Ribosomal Database Project v. 11.5 inferred taxonomy of the 16S rRNA gene Sanger clone libraries prepared by Burgess *et al*. ^9^, the Kaiju v. 1.6.2 inferred taxonomy for the Sanger metagenomes prepared by TIGR and the Kaiju v. 1.6.2 inferred taxonomy for the Solexa reads which were later assembled to bin the MAGs. The different circles represent the 16S rRNA genes (blue), the MAGs (yellow), the Sanger metagenomes (orange) and the Solexa reads (magenta).

### Comparison of genera found in both pools

Due to the diverging biogeochemistry between ARK and ZAV, we were interested in if shared genera between pools would be more similar to each other or to existing reference genomes. To investigate this question, we focused our comparisons on two genera, *Desulfurella* and *Sulfurihydrogenibium*, for which draft MAGs were obtained in both pools with high completion (> 90%).

For *Desulfurella*, the MAGs obtained from both pools at first visually appeared to be more distantly related to each other than to existing reference genomes with ZAV-10 grouping with *Desulfurella multipotens* (Fig. 4). However, when we calculated pairwise average nucleotide identities (ANI), we found that ARK-08, ZAV-10, *D. multipotens* and *D. acetivorans* should all be considered the same species (ANI > 95%; Supplementary Table S9). A threshold of greater than 95% ANI is generally considered appropriate for assigning genomes to the same species ^30^

**Figure 4.**
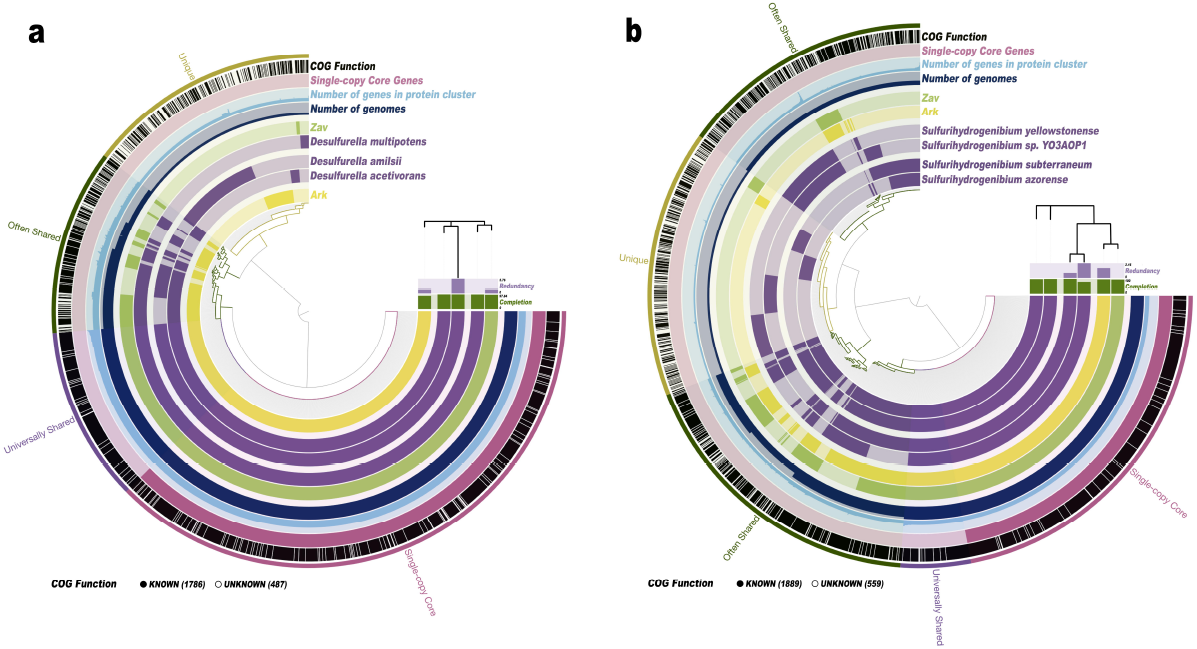
Pangenomic comparison of shared genera between pools. *Desulfurella* genera (a) and *Sulfurihydrogenibium* genera (b) identified in both Arkashin Schurf (ARK) and Zavarzin Spring (ZAV) are visualized respectively in anvi’o against reference genomes downloaded from NCBI. ARK-08 and ZAV-10 were compared to three representative *Desulfurella* genomes including *D. multipotens* (GCA_900101285.1), *D. acetivorans* (GCA_000517565.1) and *D. amilsii* (GCA_002119425.1), while bins ARK-13 and ZAV-16 were compared to four representative *Sulfurihydrogenibium* genomes including S. *subterraneum* (GCA_000619805.1), S. *yellowstonense* (GCA_000173615.1), S. *azorense* (GCA_000021545.1) and S. *sp. YO3AOP1* (GCA_000020325.1). Genomes are arranged based on a phylogenetic tree of shared single-copy core genes produced in anvi’o using FastTree v. 2.1. Protein clusters have been grouped into categories based on presence/absence including: ‘Single-copy core genes’ (protein clusters representing #genes from Campbell *et al*. ^64^), ‘Universally shared’ (protein clusters present in all genomes), ‘Often Shared’ (protein clusters present in two or more genomes) and ‘Unique’ (protein clusters present in only one genome). Gene calls were annotated in anvi’o using NCBI’s Clusters of Orthologous Groups (COG’s). Protein clusters with an assigned NCBI COG are indicated in black.

For *Sulfurihydrogenibium*, the two MAGs appear to be more closely related to each other than to existing reference genomes and appear to form their own clade (Fig. 4B). This inference is supported by high ANI values from which we infer that the two MAGs are actually the same species (ANI = 97.2%; Supplementary Table S10). The ANI values of these MAGs suggest that they comprise a distinct species for this genus when compared to the four existing reference genomes (ANI < 76%).

### Functional annotations; qualitative differences between ARK and ZAV

High concentrations of arsenic have previously been found in Arkashin Schurf, hence we searched specifically for homologs of genes involved in arsenic biotransformation and compared them between the two pools. Homologs of genes encoding proteins that are predicted to be involved in the arsenic biogeochemical cycle were present in both pools (n = 86 for ARK and n = 73 for ZAV; Supplementary Table S11). These included homologs of ArsA, ArsB, ArsC and ArsH, ACR3, Arsenite_ox_L, Arsenite_ox_S, and the ArsR regulator (Supplementary Table S12). Homologs of ArsH were restricted to Arkashin Schurf. ACR3 could be assigned to MAG ARK-10; ArsA to ARK-07, ARK-11, ARK-16, and ZAV-03; Arsenite_ox_L to ARK-07 and ARK-16; and Arsenite_ox_S to ARK-01 and ARK-07.

We searched for complete chemical pathways in both pools and linked them to our MAGs. In total, 222 complete KEGG gene pathways were predicted to be present in ARK and ZAV combined. Completeness of a pathway is defined as including all necessary components (gene blocks) to complete a metabolic cycle. Nine complete pathways were predicted to be present exclusively in ARK (Supplementary Table S13) and 14 pathways in ZAV. We grouped the 119 shared gene pathways that could be found in both pools based on KEGG orthologies into carbohydrate and lipid metabolism (n = 31; Supplementary Table S14), energy metabolism (n = 16; Table 4), and environmental information processing (n = 31; Supplementary Table S15). An exhaustive list of all predicted KEGG pathways including their raw copy number in both pools and KEGG pathway maps can be found in Supplementary Table S16.

**Table 4.**
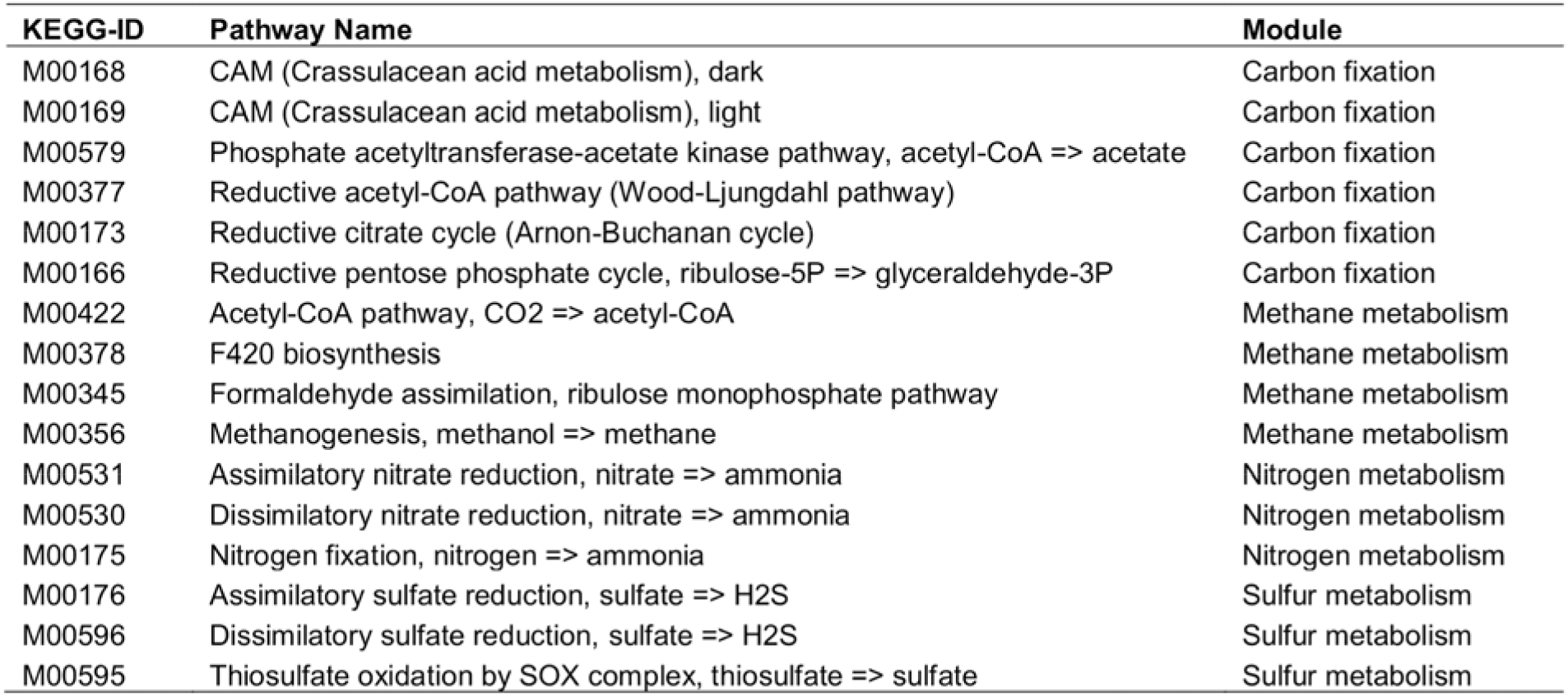
Complete energy metabolism KEGG pathways that were predicted to be present in both pools based on the recovery of putatively homologous genes. Shown are all complete KEGG pathways; *i.e*., gene pathways of which all genes (blocks) were represented (n > 5 to 1,860) in both pools (Arkashin Schurf and Zavarzin Spring). For each pathway its KEGG-ID, name and pathway module are given. Presence of pathways was predicted based on the retrieval of homologous genes.

## Discussion

We recovered 36 MAGs (20 in ZAV and 16 in ARK) comprising a broad phylogenetic range of archaeal and bacterial phyla. Moreover, the MAG’s we constructed from two volcanic hot springs expand the microbial tree of life with several new genomes from taxonomic groups that had previously not been sequenced. These include draft MAGs for several candidate phyla including archaeal Korarchaeota, Bathyarchaeota and Aciduliprofundum; and bacterial Aminicenantes. Korarchaeota and Crenarchaeota have been found in thermal ecosystems previously using 16S rRNA gene amplicon sequencing ^15,31,32^. but with very little genomic information so far ^19,33^. Korarchaeota have been described exclusively in hydrothermal environments ^34^. They belong to one of the three major supergroups in the Euryarchaeota, together with the Thaumarchaeota, Aigarchaeota and the Crenarchaeota (TACK, proposed name Eocyta^35^). TACK make up a deeply branching lineage that does not seem to belong to the main archaeal groups. Bathyarchaeota are key players in the global carbon cycle in terrestrial anoxic sediments ^36,37^. They appear to be methanogens and can conserve energy via methylotrophic methanogenesis (see below). *Aciduliprofundum* spp. have only been found in hydrothermal vents and have one cultivated representative; *Aciduliprofundum boonei* ^38^. This taxon is an obligate thermoacidophilic sulphur and iron reducing heterotroph. Aminicenantes (candidate phylum OP8) is a poorly characterized bacterial lineage that can be found in various environments, such as hydrocarbon-contaminated soils, hydrothermal vents, coral-associated, terrestrial hot springs, and groundwater samples ^39^. A high-level of intraphylum diversity with at least eight orders has been proposed for this group ^40^.

The representation of the current microbial tree of life we constructed here using 37 single-copy marker genes recapitulates and confirms the structure seen in other recent studies using different sets of single copy genes ^27,41^. The placement of the MAGs into this phylogeny was in alignment with their taxonomic assignments based on CheckM’s marker set with two exceptions ^42^.

ARK-16 clustered with Crenarchaeota in the tree but was assigned to Korarchaeota in the assignment. Only one genomic representative is currently available for the phylum Candidatus Korarchaeota (other than ZAV-18). It is possible that ARK-16 is still a member of this phyla but is also distantly related to the available genome leading to the branching pattern observed here (Fig. 2). Other possibilities include that ARK-16 represents a novel phylum of archaea that is sister to Crenarchaeota or that ARK-16 is a new group within the Crenarchaeota.

ARK-02 clustered next to *Acidobacterium* spp. (Acidobacteriales) in the tree but was assignedto the *Aminicenantes* group with CheckM. Previous phylogenomic studies have placed *Candidatus Aminicenantes* as sister to the Acidobacteriales ^43^. Thus, the placement of ARK-02 as sister to Acidobacteriales here is likely not an ambiguity and instead further supports Candidatus Aminicenantes as its proper taxonomic placement (Supplementary Fig. S1). Originally, we had included three Candidatus Aminicenantes species when building Fig. 2, but they were removed by trimAl because they were missing a substantial number of the 37 single-copy marker genes.

Generally, many of the same taxa as the MAGs assembled here were also observed in the analysis of the 16S rRNA gene sequence library prepared by Burgess *et al* ^9^. Using an updated version of the RDP database decreased the proportion of unclassified sequences in the ARK bacterial library and the ZAV archaeal library, identifying several new phyla in both sequence libraries. The proportion of unclassified sequences in the ZAV bacterial library decreased slightly but remained relatively high (17.3%; Supplementary Table S6), indicating that there is still a large amount of bacterial novelty in ZAV. It is possible that this proportion can be partially explained by the five bacterial ZAV MAGs that do not represent genera identified in the clone library, but which were observed in the Sanger metagenomic reads. These include members of the phyla Aquificae, Bacteriodetes, Thermodesulfobacteria and Thermotogales (Table 3; Supplementary Table S8).

A recent 16S rRNA gene sequencing study by Merkel *et al*. ^8^. found the most abundant members of ARK to be the archaea Thermoplasmataceae group A10 (phylum Euryarchaeota) and *Caldisphaera* at 34% and 30% relative abundance respectively. Here, we generated two medium quality draft MAGs for *Caldisphaera* (ARK-06; ARK-07). We did not find any members of the Thermoplasmataceae group A10, but we did generate a high-quality draft MAG from a candidate group in the same phylum, *Candidatus Aciduliprofundum* (ARK-15). In Fig. 2, the closest relative to ARK-15 is *Aciduliprofundum sp. MAR08-339* and together they from a sister group to *Picrophilus oshimae*, a member of the Thermoplasmataceae. *Candidatus Aciduliprofundum* has been previously placed next to Thermoplasmataceae based on a maximum likelihood tree using 16S rRNA genes ^38^ and a Bayesian phylogeny constructed from the concatenation of 57 ribosomal proteins ^44^. However, the relationship of *Aciduliprofundum* to other archaea is still unresolved ^45^.

Even though the archaeal tree of life has expanded to include more genomic representatives since the Burgess *et al*. ^9^ study, our understanding of the archaeal tree of life is still limited. Primer bias is known to historically plague archaeal amplicon 46 sequencing studies ^46^ and additionally may explain why Burgess *et al*. were unable to amplify archaeal sequences from ARK. The novel archaeal MAGs assembled here combined with the additional archaeal and bacterial phyla identified in the Sanger metagenomes provides a good argument for re-examining previously characterized environments using new methods to further expand our view of the tree of life (Fig. 3).

After investigating the relatedness of *Desulfurella* draft MAGs from both pools (ARK-08 and ZAV-10) to existing reference genomes, we propose that the MAGs, *D. multipotens* and *D. acetivorans* should all be considered the same species. The collapse of *D. multipotens* and *D. acetivorans* into one species has been previously suggested by Florentino *et al*. based on both ANI and DDH (DNA-DNA hybridization) values ^47^. Two species of *Desulfurella* were previously isolated from Kamchatka, *D. kamchatkensis* and D. *propionica* ^48^. Neither strain has been sequenced, although Miroshnichenko *et al*. performed DDH between these strains and *D. acetivorans* finding values of 40% and 55% respectively indicating that these are unique strains. However, the authors also found that there was high sequence similarity (> 99%) between full length 16S rRNA genes for *D. multipotens, D. acetivorans, D. kamchatkensis* and *D. propionica*. Given this, and that the MAGs, *D. multipotens* and *D. acetivorans* appear to be one species, we wonder what the genomes of *D. kamchatkensis* and *D. propionica* might reveal about the relationships within this genus. This situation highlights the need for an overhaul in microbial taxonomy based on whole genome sequences, a concept which has been previously discussed by Hugenholtz *et al*. ^49^ and recently been proposed in Parks *et al*. ^41^.

Meanwhile, the *Sulfurihydrogenibium* MAGs (ARK-13 and ZAV-16) appear to be from a previously unsequenced species for this genus when compared to existing reference genomes. It is possible that these MAGs represent draft genomes of S. *rodmanii*, a novel species of *Sulfurihydrogenibium* that was previously cultured from hot springs in the Uzon Caldera, but for which a reference genome does not yet exist ^13^. *Sulfurihydrogenibium rodmanii* is a strict chemolithoautotroph, it is microaerophilic and utilizes sulphur or thiosulfate as its only electron donors and oxygen as its only electron acceptor. ARK-13 has a GC content of 34.23% and ZAV-16 has a GC content of 34.32% (Table 1). These closely match the GC content estimate reported for S. *rodmanii* of 35% ^13^. Additionally, S. *rodmanii* is the best match for several of the *Sulfurihydrogenibium* sequences in the Burgess *et al*. clone library, providing further support for the hypothesis that the MAGs identified in this study may represent members of this species.

Out of the total 222 predicted complete KEGG pathways, only nine were unique to ARK and 15 to ZAV. Interestingly, homologs of the complete denitrification pathway M00319 and anoxygenic photosystem II M00597 were only found in ZAV. Denitrification is a respiration process in which nitrate or nitrite is reduced as a terminal electron acceptor under low oxygen or anoxic conditions and in which organic carbon is required as an energy source ^8^. Denitrification has been predicted to be carried out mostly by *Thiobacillus* spp., *Micrococcus* spp., *Pseudomonas* spp., *Achromobacter* spp., and *Calditerrivibrio* spp. in the Uzon Caldera ^16^. The last of these is represented here by ZAV-05 which most probably contributed homologs to this complete, predicted KEGG pathway. This is in agreement with Burgess *et al.’s* stable isotope analysis of N^15^. Anoxic photosynthesis is performed by obligate anaerobes such as *Chloroflexus* and *Roseiflexus* and requires energy in the form of sunlight. Hence, anoxic photosynthesis is most likely carried out at the surface and in the water column of ZAV by these organisms.

We reconstructed 31 different predicted carbohydrate and lipid metabolism KEGG pathways using homologous genes in ARK and ZAV including cell wall component biosynthesis; *e.g*., isoprenoids and other lipopolysaccharides. Both pools are predicted to contain genes that encode proteins for all major aerobic energy cycles, including the complete citrate cycle, Entner-Doudoroff, Leloir, and Embden-Meyerhof pathway. Carbon fixation through autotrophic CO_2_ fixation was represented by four major pathways in both pools: crassulacean acid metabolism (CAM), Wood-Ljungdahl pathway, Arnon-Buchanan cycle, and reductive pentose phosphate cycle. There are many variants of the Wood-Ljungdahl pathway, one of which is preferred by sulphate-reducing microbial organisms that grow by means of anaerobic respiration ^50^. Coupled with methanogenesis; *i.e*., the Acetyl-CoA and the F420 pathway reducing CO_2_, this represents one of the most ancient metabolisms for energy generation and carbon fixation in archaea ^51^. Recently it has been shown that Bathyarchaeota possess the archaeal Wood-Ljungdahl pathway _36,37_. Similarly, the Arnon-Buchanan cycle is commonly found in anaerobic or microaerobic microbes present at high temperatures, such as *Aquificae* and *Nitrospirae* ^52^.

We found homologs of three predicted, complete major nitrogen pathways and three complete major sulphur pathways in both pools. These included assimilatory nitrate reduction to ammonia, dissimilatory nitrate reduction to ammonia and nitrogen fixation from nitrogen to ammonia. With regard to sulphur, the metabolisms included thiosulfate oxidation to sulphate, assimilatory sulphate reduction to H2S, and dissimilatory sulphate reduction to H_2_S. These two pathways; *i.e*., aerobic sulphur oxidation and anaerobic hydrogen oxidation coupled with sulphur compound reduction can be performed by aerobic *Sulfurhydrogenibium* and anaerobic *Caldimicrobium* ^8^. We assembled MAGs of both taxa in ZAV (ZAV-16 and ZAV-15) and of the former taxon in ARK (ARK-13). Sulphate-reducing bacteria can also change the concentration of arsenic in a pool by generating hydrogen sulphide, which leads to reprecipitation of arsenic ^53^. Hence, the presence of sulphur oxidizers and sulphur reducers in a pool can significantly impact the fate of environmental arsenic.

Homologs of predicted protein families that play a role in the biotransformation of arsenic were found in both pools. Such as homologs of the predicted genes *arsB* and *ACR3*, which code for arsenite (As(III)) pumps that remove reduced arsenic from the cell ^54^. Early microorganisms originated in anoxic environments with high concentrations of reduced As(III) ^55^. Most microbes have evolved efflux systems to get rid of As(III) from their cells ^54^. Hence, nearly every extant microbe is armed with As(III) permeases, such as ArsB or ACR3 ^53^. Some organisms evolved genes encoding anaerobic respiratory pathways utilizing As(III) as an electron donor to produce energy while oxidizing As(III) to As(V) ^56^. This type of arsenic cycling has been predicted to be carried out by members of *Hydrogenobaculum* spp., *Sulfurihydrogenibium* spp., *Hydrogenobacter* spp., and other Aquificales ^57^. We found MAGs of *Hydrogenobaculum* and *Sulfurihydrogenibium* present in both pools, with two high quality drafts in ARK. In addition to the Aquificales, we also found several copies of ACR3 in ARK-10, *Caldisericum exile*.

With increasing atmospheric oxygen concentrations, As(III) is oxidized to As(V), a toxic compound which can enter the cells of most organisms via the phosphate uptake systems ^54^. Consequently, organisms needed to find ways to survive with these environmental toxins inside their cells. This was the advent of several independently evolved As(V) reductases, such as the *arsC* system ^53^. The only protein homolog which was found exclusively in ARK is ArsH, which is presumably involved in arsenic methylation. In addition to oxidation and reduction of inorganic arsenic species, arsenic methylation is another strategy to detoxify As(V) ^53^. Common methylation pathways include ArsM and ArsH and are regulated by the As(III)-responsive transcriptional repressor ArsR, which was common in both pools. Coupled with ATP hydrolysis, some microbes also developed an energy-dependent process where As(III) is actively pumped out of the cell ^53,58^. Driven by the membrane potential, ArsA can bind to ArsB and pump out As(III). Homologs of predicted ArsB proteins were found in both pools with no assignment to any of our MAGs. However, homologs of *arsA* could be found in ARK-07, ARK-11, ARK-16, and ZAV-03 which correspond to a Korarchaeota representative, a *Caldispaera* sp. and two *Mesoaciditoga* species, one in each pool.

The breadth of undiscovered microbial diversity on this planet is extreme, particularly when it comes to uncultured archaea whose abundance has likely been underestimated because of primer bias for years ^46^. By incorporating metagenomic computational methods into the wealth of pre-existing knowledge about these ecosystems, we can begin to putatively characterize the ecological roles of microorganisms in these hydrothermal systems. Future work should aim to isolate and characterize the novel microorganisms in these pools so that we can fully understand their biology, confirm the ecological roles they play, and complement and expand the current tree of life.

## Methods

### Sample collection and DNA extraction

The DNA used here is the same DNA that was used in Burgess *et al*. ^9^. In short, Burgess *et al*. extracted DNA from sediment from ARK (Fig. 1), collected in the field in 2004 (sample A04) using the Ultra-Clean ^®^ Soil DNA Kit (MoBio Laboratories, Inc., Carlsbad, CA, USA) following the manufacturer’s instructions. They then extracted DNA from sediment from ZAV (Fig. 1) collected in the field in 2005 (sample Z05) using the PowerMax ^®^. Soil DNA Isolation Kit (MoBio Laboratories, Inc.) following the manufacturer‘s instructions. DNA was sequenced with two approaches: Sanger sequencing of clone libraries at TIGR (The Institute for Genomic Research) and paired-end Solexa3 sequencing of 84 bp at the UC Davis Genome Center. Details on sequencing and clone library construction can be found in the Supplementary Material.

### Sequence processing and metagenomic assembly

Quality filtering was performed on Solexa reads using bbMap v. 36.99 ^59^ with the following parameters: qtrim = rl, trimq = 10, minlength = 70; *i.e*., trimming was applied to both sides of the reads, trimming the reads back to a Q10 quality score and only keeping reads with a minimum length of 70 bp. Adaptors were removed from the Solexa reads and samples were assembled into two metagenomes (one for ARK and one for ZAV) using SPAdes v. 3.9.0 ^60^ with default parameters for the metagenome tool (metaspades). Sanger metagenomic reads were processed using phred ^61^ to make 62 base calls and assign quality scores, and Lucy ^62^ to trim vector and low quality sequence regions.

### Metagenomic binning and gene calling

Metagenomic data was binned using anvi’o v. 2.4.0 ^29^, following a modified version of the workflow described by Eren *et al. ^29^*. First, a contigs database was generated for each sample from the assembled metagenomic data using ‘anvi-gen-contigs-database’ which calls open reading frames using Prodigal v. 2.6.2 ^63^ Single copy bacterial ^64^. and archaeal ^65^ genes were identified using HMMER v. 3.1b2 ^66^. Taxonomy was assigned to ^22^. contigs using Kaiju v. 1.5.0 ^22^ with the NCBI BLAST non-redundant protein database *nr* including fungi and microbial eukaryotes v. 2017–05–16. In order to visualize the metagenomic data with the anvi’o interactive interface, a blank-profile for each sample was constructed with contigs > 1 kbp using ‘anvi-profile’, which hierarchically clusters contigs based on their tetra-nucleotide frequency profiles. Contigs were manually clustered into bins using a combination of hierarchical clustering, taxonomic identity, and GC content using ‘anvi-interactive’ to run the anvi’o interactive interface. Clusters were then manually refined using ‘anvi-refine’ and bins were continuously assessed for completeness and contamination using ‘anvi-summarize’ and the CheckM v. 1.0.7 ^42^ lineage-specific workflow. For a detailed walk-through of the analyses used to bin the metagenomic data, please refer to the associated Jupyter notebooks for ZAV ^67^ and for ARK^68^.

### Taxonomic and phylogenetic inference of MAGs

Using the standards suggested by Bowers *et al*. ^69^, bins were defined as high-quality draft (> 90% complete, < 5% contamination), medium-quality draft (> 50% complete, < 10% contamination) or low-quality draft (< 50% complete, < 10% contamination) MAGs.

Taxonomy was tentatively assigned to MAGs using a combination of inferences by Kaiju ^22^ and CheckM’s lineage-specific workflows ^42^. Taxonomy was refined and confirmed by placing MAGs in a phylogenetic context using PhyloSift^70^ v. 1.0.1 with the updated PhyloSift markers database (version 4, 2018–02–12 ^71^). For this purpose, MAGs, all taxa previously identified by Burgess *et al*. ^9^ with complete genomes available on NCBI (downloaded 2017–09–06), and all archaeal and bacterial genomes previously used in Hug *et al*. (2016) were placed in a phylogenetic tree ^27^. Details on how this tree was constructed can be found in the Supplementary Material. Briefly, PhyloSift builds an alignment of the concatenated sequences for a set of core marker genes for each taxon. We used 37 of these single-copy marker genes (Supplementary Material) to build an amino acid alignment, which was trimmed using trimAl v.1.2 ^72^. Columns with gaps in more than 5% of the sequences were removed, as well as taxa with less than 75%of the concatenated sequences. The final alignment ^73^ comprised 3,240 taxa (Supplementary Table S3) and 5,459 amino acid positions. This alignment was then used to build a new phylogenetic tree in RAxML v. 8.2.10 on the CIPRES Science Gateway web server ^74^ with the LG plus CAT (after Le and Gascuel) ^75^ AA substitution model. One hundred fifty bootstrap replicates were conducted. The full tree inference required 2,236 computational hours on the CIPRES supercomputer. The Interactive Tree Of Life website iTOL was used to finalize and polish the tree for publication ^76^. All genomes used in this tree and a mapping file can be found on Figshare (genomes in Hug *et al.’s* tree of life (2016) ^77–79^ and genomes from Burgess *et al*. (2012) ^80^).

### New analysis of 16S rRNA gene sequences

The 16S rRNA gene sequences generated by Burgess *et al*.^9^ were downloaded from NCBI. Using the same parameters and method as described in Burgess *et al*., but with an updated database, we inferred the taxonomy of these sequences. Briefly, sequences were uploaded to the RDP (Ribosomal Database Project) website and aligned to the RDP database (v. 11.5) ^81^. Then, the SeqMatch tool was used to identify the closest match using all good quality sequences ≥ 1200 bp in length.

### Taxonomic inference of metagenomic reads

Taxonomy was assigned to the quality-filtered Solexa reads for each sample using Kaiju v. 1.6.2 ^22^ with the NCBI BLAST non-redundant protein database *nr* including fungi and microbial eukaryotes v. 2017-05-16. Kaiju was run using greedy mode with five substitutions allowed with an e-value cut-off for taxonomic assignment of 0.05. Taxonomy for each sample was summarized by collapsing taxonomic assignments to the phylum level. This process was repeated to infer taxonomy for the metagenomic reads from the Sanger clone libraries. Inferred taxonomy for the Solexa reads, Sanger metagenomes, MAGs and the RDP results for the 16S rRNA genes sequences from Burgess *et al*. were then imported into R v. 3.4.3 and compared using the ‘VennDiagram’ package v.1.6.20 ^82^.

### Pangenomic comparison of pools and investigation of arsenic metabolizing genes

In order to characterize gene functions in ARK and ZAV, we identified protein clusters within the two thermal pools and visualized them in anvi’o using their pangenomic workflow ^83^. We also used this workflow to investigate whether shared genera between pools would be more similar to each other or to reference genomes. We focused our comparisons on the genera *Desulfurella* and *Sulfurihydrogenibium* as we were able to obtain draft MAGs for these genera in both pools with high completion (> 90%). Bins ARK-08 and ZAV-10 were compared to all three representative *Desulfurella* genomes available on NCBI (GCA_900101285.1, GCA_000517565.1, and GCA_002119425.1), while bins ARK-13 and ZAV-16 were compared to all four representative *Sulfurihydrogenibium* genomes (GCA_000619805.1, GCA_000173615.1, GCA_000021545.1, and GCA_000020325.1).

To quantify pairwise similarities, we used DIAMOND v. 0.9.9.110 ^84^, which calculates similarities between proteins. Then, we applied the MCL algorithm v. 14-137 ^85^ to construct protein clusters with an inflation value of 2.0 when comparing all MAGs from both pools and 10.0 when investigating close relatives, and muscle v. 3.8.1551 to align protein sequences ^86^. Gene calls were annotated during this workflow with NCBI’s Clusters of Orthologous Groups (COGs ^87^) and Kyoto Encyclopedia of Genes and Genomes (KEGG) orthologies downloaded from GhostKOALA ^88^ following the workflow for anvi’o by Elaina Graham (as described in http://merenlab.org/2018/01/17/importing-ghostkoala-annotations/).

Given unusually high concentrations of arsenic in Arkashin Schurf, we decided to look specifically for homologs of genes involved in arsenic biotransformations and compare them between the two pools. We manually downloaded HMMs (Hidden Markov Models) for protein families with a functional connection to arsenic from the TIGRFAM repository. Our selection of proteins was based on ^Zhu *et al*. (2017)^ ^53^ (i.e., ArsA, ArsB, ArsC, ArsH, ArsR, and ACR3). We searched all open reading frames (ORFs) in both pools against them using blastx v. 2.6.0. Hits with at least 85% coverage on the length of the match, an e-value of 1e-10 and 85% identity were kept. These hits were searched using blastx v. 2.6.0 with an e-value threshold of 1e-4 against the MAGs to find out which organisms possess genes involved in arsenic biotransformations ^89^.

R v. 3.4.0 with the package ‘plyr’ v. 1.8.4 ^90^ was used to summarize homologs of shared and unique genes and predicted metabolic pathways qualitatively between the two pools. When investigating close relatives, phylogenetic trees were built in anvi’o using FastTree ^91^ on the single-copy core genes identified to order taxa during visualization. Average nucleotide identity (ANI) values were calculated between close relatives and representative genomes using autoANI (https://github.com/osuchanglab/autoANI; ^30,92–94^). Adobe Photoshop CS6 was used to finalize figures.

### Data availability statement

Sanger reads were deposited on NCBI’s GenBank under SRA IDs SRS3441489 (SRX4275258) and SRS3441490 (SRX4275259). Raw Solexa reads were deposited on NCBI’s GenBank under BioProject ID PRJNA419931 and BioSample IDs SAMN08105301 and SAMN08105287; *i.e*., SRA IDs SRS2733204 (SRX3442520) and SRS2733205 (SRX3442521). Draft MAGs were deposited in GenBank under accession numbers SAMN08107294 - SAMN08107329 (BioProject ID PRJNA419931). NCBI performed their Foreign Contamination Screen and removed residual sequencing adaptors prior to publication. Draft MAGs and associated anvi’o files can be found on DASH ^95^. The alignment and raw tree file in Newick format used for Fig. 2 can be found on Figshare ^73,96^.

## Acknowledgments

We would like to thank Russell Neches (ORCID: 0000-0002-2055-8381) for the use of photographs taken on a trip to Kamchatka, Russia in 2012. We would like to thank Elizabeth A. Burgess and Juergen Wiegel for providing JAE with the DNA used here. We are also grateful to Christopher Brown (ORCID: 0000-0002-7758-6447) and Laura Hug (ORCID: 0000-0001-5974-9428) for their help getting access to the genomes used in Fig. 2. A special thank you goes to Alexandros Stamatakis (ORCID: 0000-0003-0353-0691), Wayne Pfeiffer and Mark Miller for offering their help with getting the CIPRES Science Gateway server to run RAxML-HPC2 v.8 on XSEDE.

## Author Contributions

LGEW and CLE analysed the data, prepared figures and/or tables, wrote, edited and reviewed drafts of the paper. GJ advised on data analysis and reviewed drafts of the paper. JAE contributed reagents/materials/analysis tools, advised on data analysis, reviewed and edited drafts of the paper.

## Additional Information and Declarations

### ORCIDS

LGEW: orcid.org/0000-0003-3632-2063

CLE: orcid.org/0000-0001-7334-403X

GJ: orcid.org/0000-0002-8746-2632

JAE: orcid.org/0000-0002-0159-2197

